# Durability of DNA-LNP and mRNA-LNP Vaccine-Induced Immunity Against SARS-CoV-2 XBB.1.5

**DOI:** 10.1101/2025.10.15.682549

**Authors:** Levi Tamming, Casey Lansdell, Wanyue Zhang, Diana Duque, Jegarubee Bavananthasivam, Grant Frahm, Annabelle Pfeifle, Sathya N. Thulasi Raman, Jianguo Wu, Caroline Gravel, Andrew Stalker, Matthew Stuible, Yves Durocher, Wangxue Chen, Lisheng Wang, Simon Sauve, Anh Tran, Michael J.W. Johnston, Xuguang Li

## Abstract

mRNA-lipid nanoparticle (LNP) vaccines induce robust adaptive immune responses and have proven highly effective against SARS-CoV-2. However, their long-term effectiveness is limited by waning humoral responses, which decline substantially within the first six months post-boost vaccination. DNA-LNPs are being investigated as an alternative vaccine platform, offering prolonged antigen expression and robust immunity. Here, we present the first comparison of SARS-CoV-2 DNA- and mRNA-LNP vaccines in a long-term in vivo challenge model. Both nucleic acid platforms induced strong neutralizing antibody responses and conferred equivalent protection in Syrian hamsters challenged three weeks post-boost. Notably, DNA-LNP vaccination maintained high binding and neutralizing antibody titers six months post-boost, whereas mRNA-LNPs exhibited a marked decline. Correspondingly, while DNA-LNPs completely protected from weight loss, viral replication, and lung pathology at this late timepoint, mRNA-LNP vaccination conferred minimal protection. These findings demonstrate that DNA-LNPs can sustain durable immunity, highlighting their potential as a next-generation vaccine platform that could reduce the need for frequent boosters.

## Introduction

Ideal vaccines elicit durable immunity against their target pathogen. While mRNA vaccines have been highly effective against COVID-19, their ability to prevent symptomatic infection wanes dramatically over time ^1–6^. In particular, mRNA-induced antibody responses undergo an initial phase of rapid decline before stabilizing around 7-9 months post-vaccination ^7–10^. Although mRNA vaccination drives robust germinal center activity and affinity maturation ^11^, studies show it generates only limited long-lived plasma cell (LLPC) populations in the bone marrow ^12^. The decay of circulating antibody titers, combined with the emergence of immune-escaping variants, has necessitated repeated booster immunizations, often on an annual basis, to sustain a high level of efficacy ^13–15^.

Advances in lipid nanoparticle (LNP) technology have been critical to the success of mRNA vaccines, both by protecting the mRNA and facilitating its cellular uptake ^16,17^. The ionizable lipids within these formulations also exhibit immunostimulatory properties, functioning as self-adjuvants ^18,19^. Recent research has highlighted how these same LNP formulations can be adapted for DNA-based vaccination ^20–24^, DNA- and mRNA-LNPs display distinct antigen expression kinetics and modes of innate immune activation, differences that can profoundly shape the adaptive response ^21,25,26^.

Here, we directly compare the immunogenicity and long-term efficacy of DNA- and mRNA-LNP vaccines against the SARS-CoV-2 XBB.1.5 variant. XBB.1.5 became globally dominant in early 2023 and was selected as the antigen for the FDA-approved monovalent mRNA-LNP vaccines deployed during the 2023–2024 season ^27^. We previously demonstrated that fusion of CD40 ligand (CD40L) to the SARS-CoV-2 Spike (S) enhances the immunogenicity and efficacy of DNA-based vaccines ^28,29^. Using this same design, both DNA- and mRNA-LNP vaccines elicited robust initial immune responses that conferred complete protection against XBB.1.5 in Syrian hamsters. However, only the DNA-LNP vaccine provided durable immunity, sustaining high levels of binding and neutralizing antibodies and affording near-complete protection six months post-boost. These findings highlight the potential of DNA-LNP vaccines to confer lasting protection against emerging SARS-CoV-2 variants and emphasizes the importance of further exploring this platform.

## Results

### CD40L augments humoral and CD4⁺ T Cell responses induced by mRNA-LNPs in mice

Building on our previous finding that CD40L enhanced the immunogenicity of a DNA-based SARS-CoV-2 S vaccine ^28^, we first evaluated whether this strategy could similarly augment immune responses induced by mRNA-LNP vaccination. Using our established design framework, mRNAs encoding trimerized soluble XBB.1.5 Spike either alone (S_XBB.1.5_) or fused to the CD40L ectodomain (S_XBB.1.5_-CD40L) were synthesized. BALB/c mice were immunized intramuscularly with 1.25 µg of the mRNA-LNP vaccines on days 0 and 28 before being sacrificed on day 49 for immunological analysis (**Figure 1A**). Control animals were immunized with non-coding plasmid DNA (pDNA) encapsulated in the same LNP formulation, which has previously been shown not to elicit antigen-specific responses ^20,29^. Both mRNA vaccines induced robust serum anti-XBB.1.5 Spike IgG responses compared to control mice (**Figure 1B**). Notably, the S_XBB.1.5_-CD40L mRNA-LNP vaccine generated higher binding antibody levels than the non-adjuvanted vaccine. In addition, S_XBB.1.5_-CD40L induced greater IgG2a responses (**Figure 1C**) without a corresponding rise in IgG1 levels (**Figure 1D**), indicative of Th1 polarization. Interestingly, both the S_XBB.1.5_ and S_XBB.1.5_-CD40L vaccines raised comparable levels of neutralizing antibodies (NAbs) against XBB.1.5 (**Figure 1E**). Both S_XBB.1.5_ and S_XBB.1.5-CD40L_ elicited significant interferon-γ (IFNγ)-secreting splenocyte responses (**Figure 1F**). Notably, S_XBB.1.5_-CD40L induced a significantly greater number of IFNγ-secreting cells than S_XBB.1.5_. The inclusion of CD40L also increased the expression of IFNγ, TNFα, and IL-2 in CD4⁺ T cells (**Figure 1G-I**), with no significant effect on CD8⁺ T cell responses (**Figure 1J-L**). Together, these findings demonstrate that incorporation of CD40L into mRNA-based SARS-CoV-2 Spike vaccines enhances Th1-biased humoral and CD4⁺ T cell responses.

**Figure 1.**
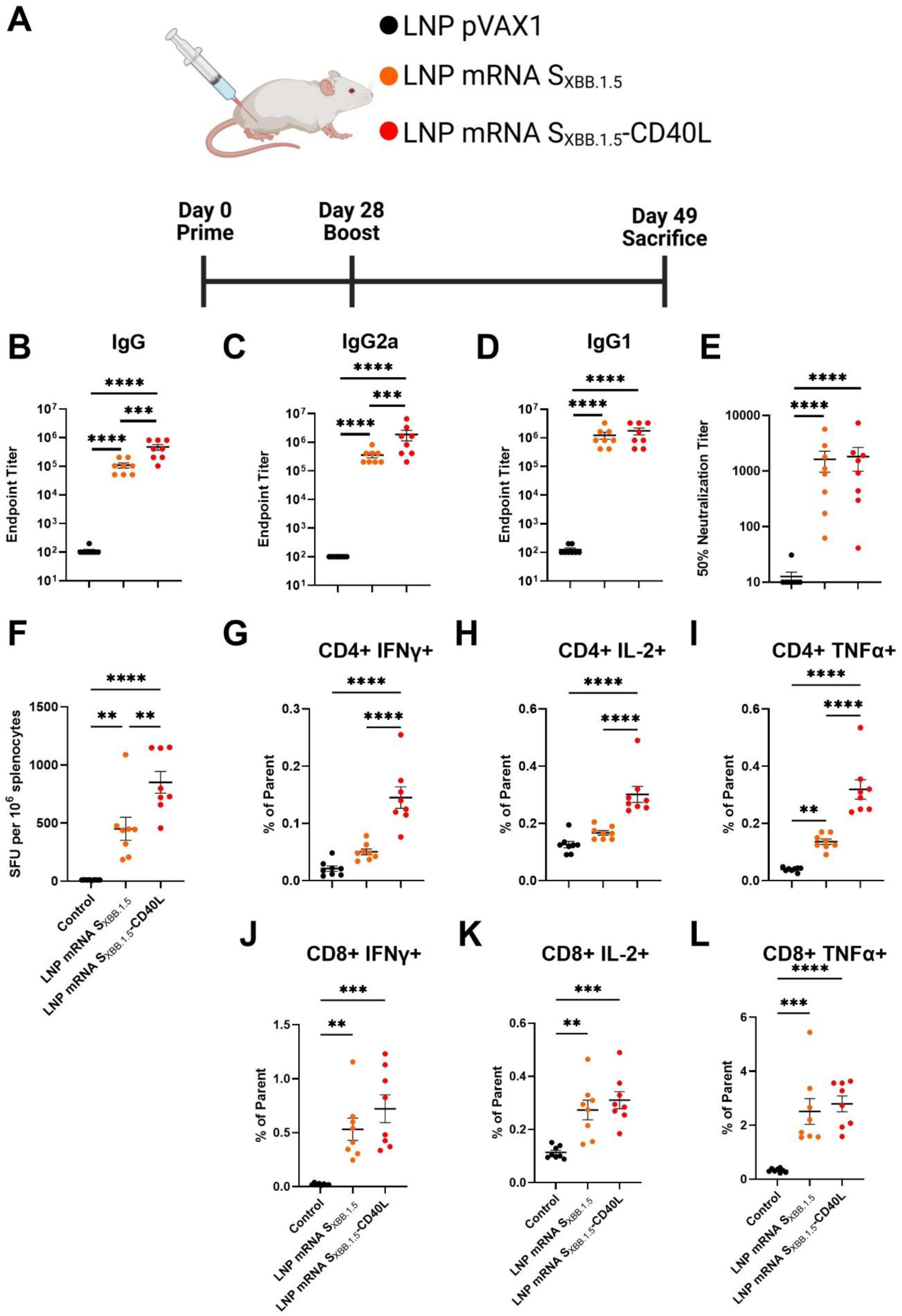
CD40L enhances immunogenicity of XBB.1.5 mRNA-LNP vaccines in BALB/c mice. **(A)** Female BALB/c mice (n = 8 per group) were immunized intramuscularly on days 0 and 28 with 1.25 µg of S_XBB.1.5_ or S_XBB.1.5_-CD40L mRNA-LNPs. Control mice were vaccinated with 2.5 µg of a non-coding pVAX1 DNA-LNP. Mice were sacrificed on day 49 to assess adaptive immune responses. XBB.1.5 Spike-specific IgG **(B)**, IgG2a **(C)** and IgG1 **(D)** levels in the sera measured by ELISA. The 50% neutralizing titer (NT50) of mouse sera was determined using XBB.1.5 pseudotyped-VSV. Cellular immune responses were assessed by stimulating splenocytes with 1 µg/mL of an XBB.1.5 Spike overlapping peptide pool. (**F**) IFNγ-secreting splenocytes as determined by ELISpot. Frequency of cytokine expression by CD4^+^ (**G-I)** and CD8^+^ (**J-L**) T cells as determined by intracellular cytokine staining. Data shown are mean ± SEM. ∗∗p < 0.01, ∗∗∗p < 0.001, ∗∗∗∗p < 0.0001.

### DNA-LNP vaccines induce long-lived humoral immunity in Syrian hamsters

Given the limited durability of humoral responses observed with mRNA vaccines, we sought to assess the longevity of immunity induced by CD40L-adjuvanted DNA- and mRNA-LNP vaccines. To evaluate both immunogenicity and protection, Syrian hamsters were immunized on days 0 and 28 with either 2.5 µg of mRNA S_XBB.1.5_-CD40L or 5 µg of pVAX1 S_XBB.1.5_-CD40L, each formulated with the same SM-102-based LNPs (**Figure 2A**). The 5 µg DNA-LNP dose was previously established to be effective against B.1.617.2 and BA.5 challenge ^29^. The lower 2.5 µg mRNA dose was selected based on the higher antigen expression and immunogenicity previously observed for mRNA-LNP vaccines ^25^. As a negative control, animals received 5 µg non-coding pVAX1 DNA-LNP. Hamsters were subsequently challenged either on day 49 or day 203, six months post-boost.

**Figure 2.**
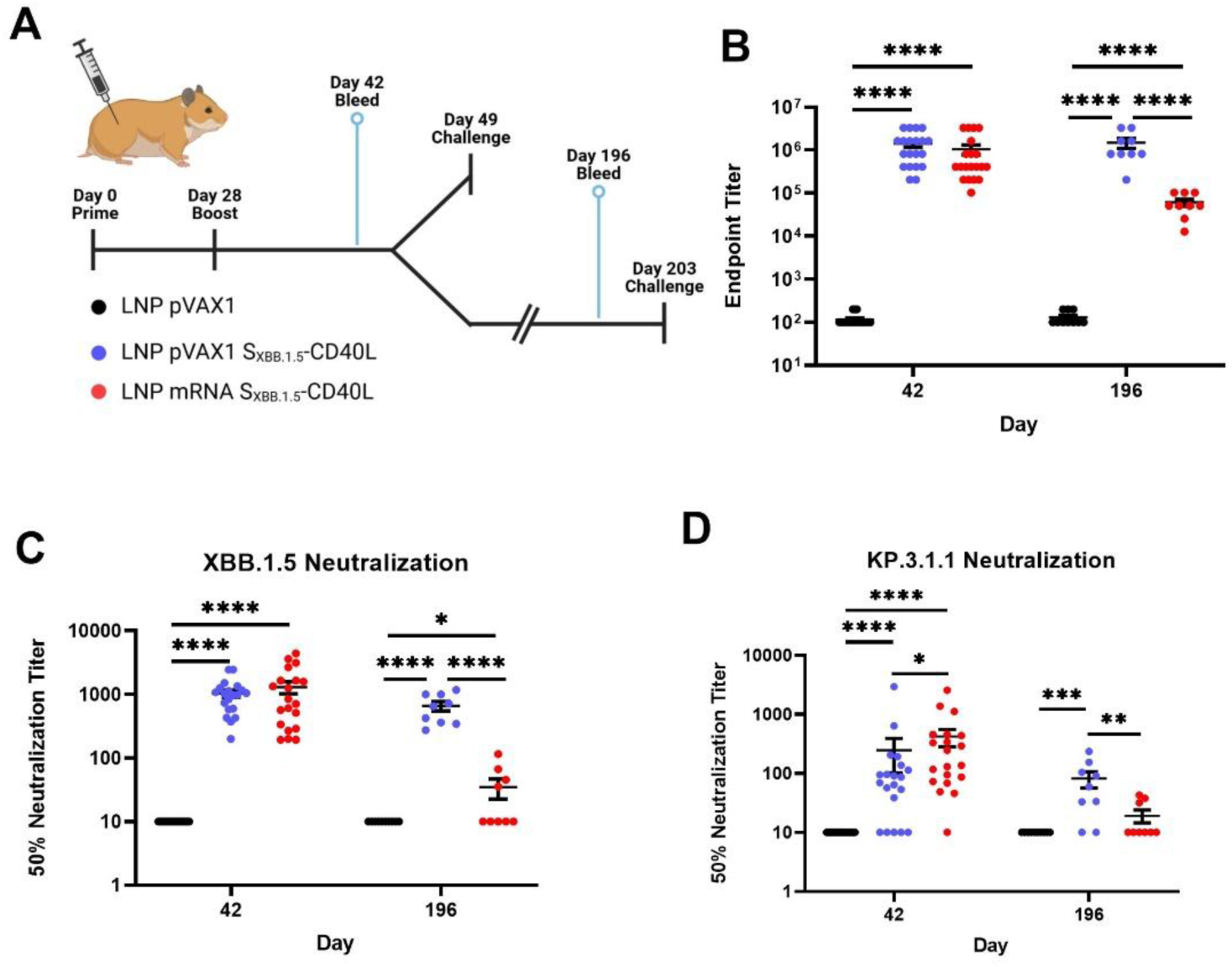
S_XBB.1.5_-CD40L DNA-LNPs induce robust and durable humoral responses in Syrian hamsters. **(A)** Male Syrian hamsters were immunized intramuscularly on day 0 and 28 with 5 µg of S_XBB.1.5_-CD40L DNA-LNP or 2.5 µg of S_XBB.1.5_-CD40L mRNA-LNP. Control hamsters were vaccinated with 5 µg of pVAX1 DNA-LNP. Animals were challenged intranasally with 1.67 × 10^5^ TCID50 of a SARS-CoV-2 XBB.1.5 isolate on day 49 or 203. Serum was collected on day 42 (n = 20 per group) or day 196 (n = 9-10 per group) to assess humoral responses. **(B)** XBB.1.5 Spike-specific IgG determined by ELISA. Serum NT50 determined using **(C)** XBB.1.5 pseudotyped-VSV or **(D)** KP.3.1.1. Data shown are mean ± SEM. ∗p < 0.05, ∗∗p < 0.01, ∗∗∗p < 0.001, ∗∗∗∗p < 0.0001.

Humoral responses in the sera were assessed at days 42 and 196. On day 42, both the DNA- and mRNA-LNP vaccines elicited robust XBB.1.5 Spike-specific IgG responses (**Figure 2B**). While binding antibody titers in the mRNA-LNP group had declined more than 10-fold by day 196, DNA-LNP vaccinated hamsters maintained high binding antibody levels. Similarly, although both vaccines elicited comparable NAb responses against XBB.1.5 at day 42, the neutralizing activity of the mRNA-LNP group was significantly diminished relative to the DNA-LNP six months post-boost (**Figure 2C**). To assess the breadth of immunity induced by both vaccines, we assessed the NAb response against KP.3.1.1 (**Figure 2D**), a more recent Omicron sublineage that demonstrated partial escape from XBB.1.5 vaccine-induced antibodies ^30^. At day 42, both DNA-LNP and mRNA-LNP immune sera neutralized KP.3.1.1, though the NAb titers were reduced compared to XBB.1.5. By day 196, neutralizing activity in the mRNA-LNP group declined to near-background levels, whereas DNA-LNP vaccinated animals retained measurable activity. Notably, immunization with DNA-LNPs encoding S_XBB.1.5_ without CD40L also induced durable humoral responses, with no apparent drop in binding antibody titers six months post-boost (**Supplementary Figure 1**). Altogether, these results highlight the capacity of DNA-LNP vaccines to elicit durable and long-lived humoral immunity against SARS-CoV-2 variants.

### DNA-LNPs provide long-term protection against XBB.1.5 SARS-CoV-2 challenge

Next, we evaluated the protective efficacy of the DNA- and mRNA-LNP S_XBB.1.5_-CD40L vaccines. Vaccinated Syrian hamsters were challenged with a matched SARS-CoV-2 XBB.1.5 isolate on day 49, three weeks post-boost vaccination (**Figure 3A**). The two S_XBB.1.5_-CD40L vaccines conferred comparable protection against weight loss following infection (**Figure 3B**). In addition, both vaccines were highly effective at suppressing viral replication in both the upper and lower respiratory tracts, with little or no infectious particles detected in either the lungs or nasal turbinates at three days post-infection (dpi) (**Figure 3C**). Consistently, subgenomic viral mRNA (sgmRNA) levels were also reduced at both sites 3-dpi (**Figure 3D**). Histopathological analysis revealed no overt signs of inflammation or infection-associated tissue damage in the lungs collected from vaccinated animals at either 3-or 6-dpi (**Figure 3E, Supplementary Figure 2**). Collectively, these data show that DNA-LNP vaccination can provide a level of protection against SARS-CoV-2 challenge that is comparable with mRNA-LNP vaccination three weeks post-boost.

**Figure 3.**
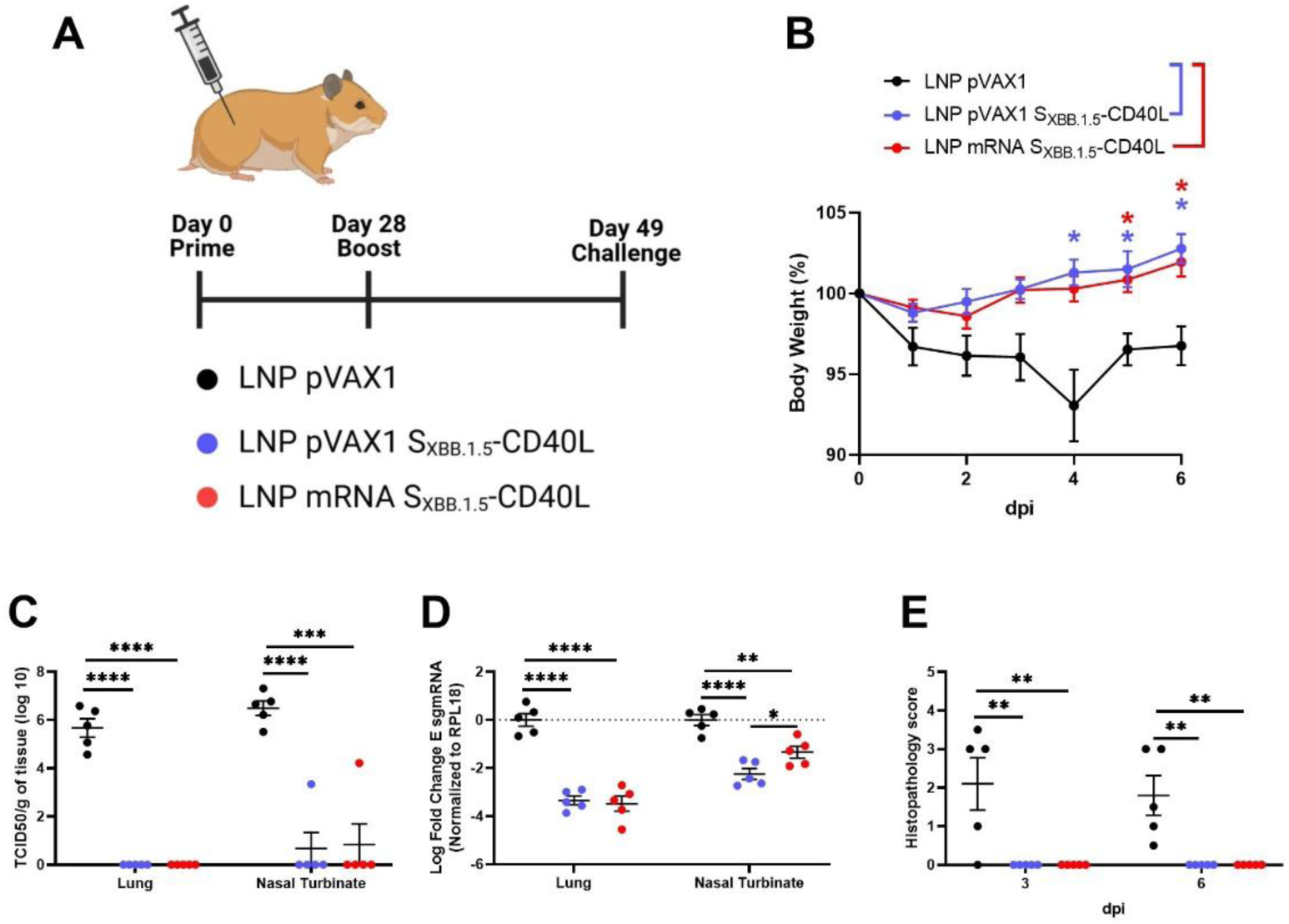
DNA- and mRNA-LNPs provide significant protection against XBB.1.5 challenge in Syrian hamsters. **(A)** Immunized Syrian hamsters were challenged intranasally with 1.67 × 10^5^ TCID50 of a SARS-CoV-2 XBB.1.5 isolate on day 49 post-vaccination and euthanized 3-(n = 5 per group) or 6-dpi (n = 5 per group). **(B)** Body weight measured daily post-challenge. **(C)** TCID50 determined in lung and nasal turbinate tissues collected 3-dpi. **(D)** SARS-CoV-2 E sgmRNA expression in the lung and nasal turbinates collected 3-dpi determined via RT-qPCR. Normalized to *RPL19* mRNA expression. **(E)** Summary of pulmonary histopathological scores 3- and 6-dpi. Data shown are mean ± SEM. ∗∗p < 0.01, ∗∗∗p < 0.001, ∗∗∗∗p < 0.0001.

To assess the durability of protection, immunized hamsters were challenged with a SARS-CoV-2 XBB.1.5 isolate on day 203, approximately six months post-boost vaccination (**Figure 4A**). While the DNA-LNP continued to afford a high level of protection at this timepoint, the mRNA-LNP provided minimal protection, mirroring the observed decline in humoral immunity. Only the DNA-LNP vaccine significantly protected against weight loss following challenge (**Figure 4B**). Correspondingly, little or no infectious virus was detected in the lungs and nasal turbinates of DNA-LNP-immunized animals at 3-dpi (**Figure 4C**). In contrast, mRNA-LNP-vaccinated animals exhibited substantially higher viral burden (**Figures 4C)** and viral sgmRNA levels (**Figure 4D**). While no overt histopathological signs of infection were detected in DNA-LNP-vaccinated hamsters at both 3- and 6-dpi, lungs from mRNA-LNP–vaccinated hamsters exhibited pronounced inflammatory cell infiltration and tissue consolidation 6-dpi (**Figure 4E, Supplementary Figure 3**). Together, these findings demonstrate that DNA-LNP vaccination sustains robust protective efficacy for at least six months post-boost vaccination.

**Figure 4.**
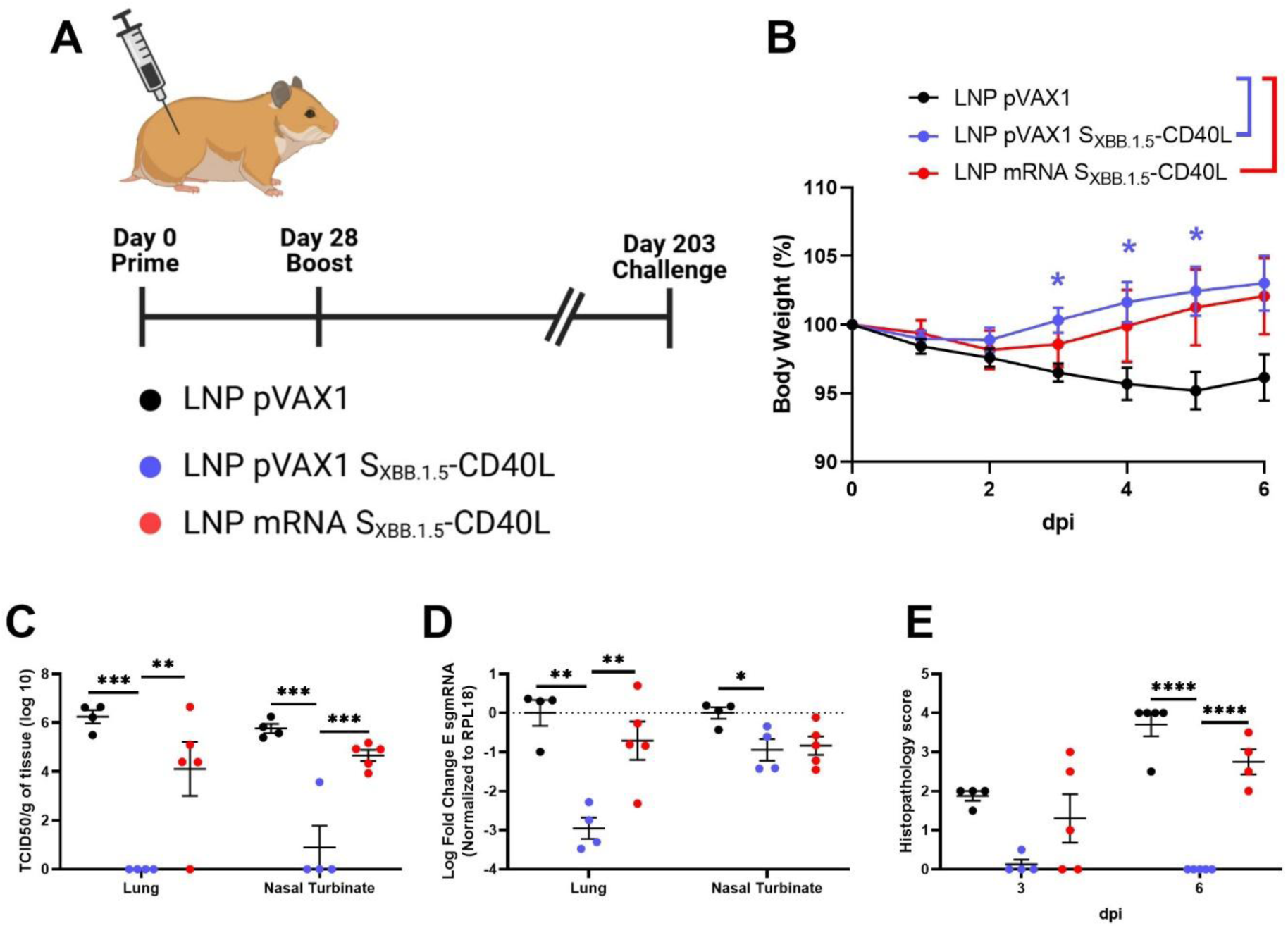
S_XBB.1.5_-CD40L DNA-LNPs provide durable protection 6-months post-boost in Syrian hamsters. **(A)** Immunized Syrian hamsters were challenged intranasally with 1.67 × 10^5^ TCID50 of a SARS-CoV-2 XBB.1.5 isolate on day 203 post-vaccination and euthanized 3-(n = 4-5 per group) or 6-dpi (n = 4-5 per group). **(B)** Body weight measured daily post-challenge. **(C)** TCID50 determined in lung and nasal turbinate tissues collected 3-dpi. **(D)** SARS-CoV-2 E sgmRNA expression in the lung and nasal turbinates collected 3-dpi determined via RT-qPCR. Normalized to *RPL19* mRNA expression. **(E)** Summary of pulmonary histopathological scores 3- and 6-dpi. Data shown are mean ± SEM. ∗p < 0.05, ∗∗p < 0.01, ∗∗∗p < 0.001, ∗∗∗∗p < 0.0001.

## Discussion

In this study, we demonstrate that DNA-LNP vaccines provide durable immunity against the SARS-CoV-2 XBB.1.5 variant, affording better long-term protection than a similar mRNA-LNP formulation. Recent studies have highlighted DNA-LNPs as a promising next-generation vaccine platform ^20–23,31–34^. Some of these studies have demonstrated that DNA-LNPs are capable of inducing comparable or even superior immune responses than mRNA-LNPs ^21,22^. Notably, these studies have also highlighted how DNA- and mRNA-LNPs differ in terms of their antigen kinetics and immune engagement. Whereas mRNA-LNPs generate potent but short-lived antigen expression, DNA-LNPs sustain expression for much longer, with detectable expression often persisting beyond one month ^21,25,26^. DNA-LNPs also appear to be more inflammatory than nucleoside-modified mRNA-LNPs, triggering strong innate immune responses through STING-sensing pathways ^21,26^. How these factors influence the overall adaptive immune response induced by DNA-LNPs remains to be fully elucidated. While a few studies have reported the durability of humoral and cellular responses elicited by DNA-LNPs ^21,22^, none to our knowledge have directly demonstrated their ability to confer long-term protection against viral challenge.

We have previously established that fusion of CD40L to the SARS-CoV-2 S enhances the magnitude and breadth of humoral responses induced by DNA and DNA-LNP vaccines ^28,29^. As such, we first assessed whether the benefits of this approach extended to mRNA-LNP vaccines. Consistent with other platforms, S_XBB.1.5-_CD40L mRNA-LNPs induced greater humoral and CD4+ T cell responses. This enhancement appeared to primarily affect Th1 responses, which are critical for protection against intracellular pathogens such as SARS-CoV-2. To the best of our knowledge, this is the first report of an antigen-CD40L fusion delivered by mRNA-LNP vaccination against an infectious pathogen.

Prior studies investigating CD40L as a molecular adjuvant have shown its capacity to promote germinal centre (GC) formation ^35^. GC reactions are linked to the generation of high-affinity LLPCs which are the primary producers of circulating antibodies ^36^. Despite this link, the S_XBB.1.5-_CD40L mRNA-LNP vaccine did not sustain a long-lived humoral response, with antibody levels waning markedly by six months post-boost vaccination, and correspondingly afforded minimal protection at this later point. In contrast, both S_XBB.1.5_ and S_XBB.1.5_-CD40L DNA-LNP vaccines maintained robust antibody responses over the same period and preserved a high level of protection. Consistent with our observations, Tursi et al. reported that DNA-LNP induced immune responses were more durable than mRNA-LNP induced responses, having greater CD8^+^ T cell recall responses in mice one year post-vaccination and stronger humoral responses in rabbits after six months ^21^. Paradoxically, Tursi et al. also observed that DNA-LNPs generated weaker GC responses than mRNA-LNPs, suggesting that the sustained humoral immunity may be driven by other factors ^21^. The half-life of nucleoside modified mRNA upon intramuscular injection is estimated to be less than 24 h, which is reflected by its short antigen expression window ^37^. Circular RNA vaccines, which are intrinsically more resistant to degradation because of their closed-loop structure, prolong antigen expression and induce superior and more durable humoral responses ^38–40^. Likewise, mRNA-1723 readenylation by TENT5a in a subset of innate immune cells has been shown to delay transcript degradation, prolonging expression and enhancing humoral responses ^41^. These findings from mRNA vaccine studies indicate that prolonged antigen expression, whether achieved through molecular design or cellular processes, can strongly influence vaccine durability. The lower but longer antigen expression offered by DNA-LNPs may be the critical factor supporting the durability of immunity observed here and in previous studies ^21,22^. Further studies are needed to establish the full duration of antigen expression from DNA-LNPs in vivo, to delineate the cell populations sustaining this expression, and to assess whether antigen persistence alone accounts for the durability observed.

Relative to mRNA vaccines, which can immediately be translated upon endosomal escape, DNA vaccines must additionally cross the nuclear membrane to enable transcription of the encoded antigen. Even when administered at a 25-fold mass excess relative to mRNA-LNPs, DNA-LNPs induce over 10-fold less antigen expression within the first 4 weeks ^25^. Notably, most published studies comparing DNA- and mRNA-LNPs have done so using equivalent nucleic acid masses, despite this resulting in substantial differences in both the molar amounts delivered and the quantity of immunogenic lipids administered. To more fully characterize the relative strengths of each platform, future comparisons should evaluate a broad range of doses. Moreover, assessing the durability of DNA-LNP-induced immunity across multiple animal models will be critical. Liao et al. reported marked differences in the magnitude and duration of DNA- and mRNA-LNP responses across species, with DNA-LNPs eliciting the strongest immunity in hamsters ^22^. Additional studies in diverse models, genetic backgrounds and, ultimately, in humans will be essential to delineate the true potential and limitations of this platform.

In summary, this study establishes that DNA-LNP vaccines can provide durable protection against the SARS-CoV-2 XBB.1.5 variant, outperforming an otherwise comparable mRNA-LNP vaccine. We demonstrate that DNA-LNP vaccination maintains antibody responses and protective outcomes for at least six months post-boost vaccination in Syrian hamsters. These findings position DNA-LNPs as a promising next-generation vaccine modality with the potential to address the issue of rapidly declining antibody responses observed with current mRNA vaccines.

## Materials and methods

### Cell lines and viruses

BHK-21 (RRID:CVCL_1915) and HEK293T-ACE2 (SL221; RRID:CVCL_C9BE) cells were cultured in Dulbecco’s modified Eagle’s medium (DMEM, Thermo Fisher) supplemented with 25 mM HEPES, 1X non-essential amino acid (NEAA), 20 U/mL penicillin, 0.02 mg/mL streptomycin, and 10% heat-inactivated fetal bovine serum (FBS). Vero-TMPRSS2 cells (RRID:CVCL_YQ49) were cultured in DMEM with L-glutamine supplemented with 1X NEAA, 1 mM sodium pyruvate, and 5% FBS. The XBB.1.5 SARS-CoV-2 viral isolate hCoV-19/USA/MD-HP40900/2022 (NR-59104) was obtained from BEI resources. The virus was propagated and titered using Vero-TMPRSS2 cells and sequenced to confirm genetic fidelity. Passage four virus stocks were used for all experiments that required live virus.

### Animal care

Animal experiments and procedures were either approved by the Animal Care Committee at Health Canada or at National Research Council Canada (NRC). All work was performed by trained staff in accordance with regulations and guidelines set by the Canadian Council on Animal Care. All infectious work was carried out under ABSL-3 conditions at the NRC. All animals were purchased from Charles River Laboratories (Senneville, QC).

### DNA and mRNA vaccine design and synthesis

The vaccine antigens were designed as previously described ^28^. Briefly, the SARS-CoV-2 XBB.1.5 Spike (GenBank: UZG29433.1) ectodomain (residues 1-1199) was mutated to have a “GSAS” substitution at the furin cleavage site (residues 678-681) and pre-fusion stabilizing proline substitutions at residues 982 and 983. The antigen was trimerized by fusion to a T4 fibritin foldon domain. The antigen was then optionally fused to the CD40L ectodomain of Mesocricetus auratus (GenBank: XM_005084522.4, residues 118-260) or of Mus musculus (GenBank accession #NM_011616, residues 118-260). Domains were separated by flexible glycine-serine linkers sequences “GSGG”. Domain coding sequences were individually codon optimized for expression in Mus musculus and Mesocricetus auratus by GenScript. DNA coding sequences were commercially synthesized (Genscript) and then subcloned into pVAX1 (Invitrogen) using KpnI/XhoI or KpnI/NotI restriction enzymes. Bulk DNA vaccine preparations were prepared using endotoxin-free gigaprep kits (Qiagen). mRNA vaccines with matching coding sequences were commercially synthesized by TriLink Biotechnologies with a CleanCap®AG Cap 1 structure and a 120A polyadenylated tail. mRNA was DNAse and phosphatase treated and purified by silica membrane. All mRNA was fully substituted with N1-methyl-pseudouridine.

### DNA-LNP and mRNA-LNP preparation

DNA- and mRNA-LNPs were synthesized as previously described within 48 h of vaccination and stored at 4°C ^25,29^. Briefly, the aqueous phase was prepared by suspending DNA in 25 mM acetate buffer (pH 4.0) or mRNA in 50 mM citrate buffer (pH 4.0). The organic phase was prepared in ethanol and consisted of SM-102 (MedKoo Biosciences), 1,2-distearoyl-sn-glycero-3-phosphocholine (DSPC, Avanti Polar Lipids), ovine cholesterol (Cholesterol, Millipore Sigma), and DMG-PEG2000 (Millipore Sigma) at a 50:10:38.5:1.5 percent molar ratio respectively. The aqueous and organic phases were mixed with a polymer amine (N = nitrogen) group to nucleic acid phosphate (P) group (N/P) ratio of 6:1 using a NanoAssemblr Ignite instrument (Precision Nanosystems). LNPs were dialyzed against Dulbecco’s phosphate-buffered saline (DPBS) for 18 h at 4°C in a 10kDa MWCO cassette (Thermo Fisher). Prior to vaccination, LNPs were passed through a 0.22 μm filter and concentrated using an Amicon Ultra 4 10 k MWCO centrifugal concentrator (Millipore Sigma). Nanoparticle size was measured using a NanoSight NS300 (Malvern Panalytical). LNP encapsulation efficiency was measured by disrupting LNPs with 1% Triton X-100 (Millipore Sigma) and adding SYBR™ Gold dye (Thermo Fisher) (Supplementary Table 1 and 2).

### Immunization and SARS-CoV-2 challenge

All animals were randomly divided into experimental groups. Female BALB/c mice (6-8 weeks-old) were vaccinated with 2.5 µg of DNA-LNPs or 1.25 µg of mRNA-LNPs, resuspended in 50 µL of PBS. Male Syrian hamsters (6-8 weeks old) were vaccinated with 5 µg of DNA-LNPs or 2.5 µg of mRNA-LNPs, resuspended in 100 µL of PBS. Injection volumes were divided evenly between both tibialis anterior muscles with a needle syringe on day 0 and day 28. Animals were bled at the time points indicated in the figure legends/text to assess antibody responses. On day 49 or 203 post-vaccination the hamsters were intranasally challenged with 1.67 ×105 TCID50 of XBB.1.5 SARS-CoV-2. Hamsters were weighed daily and euthanized either 3-or 6-days post-challenge to assess viral burden and pathology.

### Enzyme-linked immunosorbent assay (ELISA)

Spike-specific antibody titers were determined as previously described ^28^. Briefly, Nunc MaxiSorp flat-bottom 96-well plates (Thermo Fisher) were coated with 1 μg/mL of recombinant XBB.1.5 Spike protein diluted in PBS overnight at 4°C. Recombinant XBB.1.5 Spike protein (PRO8581 [SmT2v3 (XBB.1.5)] was obtained from the National Research Council of Canada, which was produced as previously described ^42^. Plates were washed with PBS containing 0.1% Tween 20 (PBS-T) before blocking with 3% (w/v) bovine serum albumin (BSA, IgG-free) (Jackson ImmunoResearch) in PBS-T for 2 h at 37°C. After washing, plates were incubated for 1 h at 37°C with mouse or hamster serum serially diluted in blocking buffer. After additional washes, the plates were incubated with either 1:2000 HRP-conjugated anti-mouse IgG (RRID:AB_772210) or 1:4000 HRP-conjugated anti-hamster IgG (RRID:AB_2337454) for 1 h at 37°C. After a final wash, plates were developed for 5 minutes using tetramethylbenzidine (TMB) substrate (Cell Signaling Technology) before being stopped with the addition of 0.16 M sulfuric acid. Absorbance values were measured at 450 nm with a Synergy 2 (BioTek) plate reader. Endpoint titers were expressed as the reciprocals of the final detectable dilution with an optical density (OD) above the cutoff value (average OD of control samples plus three standard deviations).

### Pseudovirus neutralization assay

SARS-CoV-2 spike pseudotyped VSV was generated by concurrently infecting BHK-21 cells with G∗ΔG-VSV (Kerafast, EH1020-PM) and transfecting them pDNA vectors encoding SARS-CoV-2 S Δ19 as previously described ^43^. A pLV vector encoding XBB.1.5 was synthesized commercially by Genscript. A vector encoding KP.3.1.1 was a gift from David Nemazee (Addgene plasmid #233342). After 48 and 72 h, supernatant was collected, passed through a 0.45 μm filter and stored at −80°C until use. For neutralization assays, heat inactivated serum samples were serially diluted in a white 96-well plate and incubated with the pseudovirus for 1 h at 37°C. After, 6 × 10^4^ HEK293T-ACE2 cells were added and plates were incubated for 24 h at 37°C 5% CO_2_. Luminescence was measured using Bright-Glo luciferase reagent (Promega) and a Synergy 2 (BioTek) plate reader. 50% neutralization titers (NT50) were measured as the reciprocal dilution at which a 50% reduction in relative light units (RLU) was observed relative to no-serum controls.

### Quantification of viral burden

XBB.1.5 SARS-CoV-2 viral burden was determined as previously described ^29^. Briefly, lung and nasal turbinate tissues were homogenized in PBS using a Precellys Evolution. Spin-clarified supernatants were serially diluted before being adsorbed on Vero-TMPRSS2 cells seeded in 96-well plates for 1 h at 37°C, 5% CO_2_. After the incubation, the media was replaced and the plates were incubated at 37°C, 5% CO_2_ for 5 days. Observed cytopathic effects were recorded and the 50% Tissue Culture Infectious Dose (TCID50) was calculated per g of tissue using the Reed-Muench method ^44^.

### RNA extraction and quantitative reverse-transcription PCR (qRT-PCR)

Lung and nasal turbinate tissues were homogenized in RNA shield buffer (Zymo Research) using a Precellys Evolution. RNA was extracted from the homogenized samples using a Quick-RNA Viral Kit (Zymo Research). RNA expression was quantified using a one-step Fast Virus master mix (Thermo Fisher) and either an E sgmRNA-or RPL18-specific TaqMan primer/probe set ^45,46^. E sgmRNA expression was normalized to RPL18 expression using the ddCT method ^47^. All RT-qPCR reactions were conducted in MicroAmp Fast Optical 96 wells with an Applied Biosystems 7500 Fast Real-time PCR instrument.

### Histopathology

Histopathology processing and analysis was conducted as described previously ^28^. Briefly, right lung lobes were fixed for 72 h in 10% neutral buffered formalin and processed by routine paraffin embedding methods ^48^. Four-micrometer-thick sections were stained with hematoxylin-eosin (H&E) and examined under microscopy. The severity and extent of pneumonia was scored blinded based on previously established criteria ^49^. Representative histopathology images are shown in **Supplementary Figures 2 and 3**.

### Mouse Tissue Processing

Spleens from vaccinated mice were collected three weeks post-boost vaccination and homogenized with a gentleMACS™ Dissociator (Miltenyi Biotec) using the m_spleen_01 program. Splenocytes were filtered using a 70 µm cell-strainer before being resuspended in ACK lysis buffer for 3 minutes at room temperature. Lysis was quenched by the addition of PBS and splenocytes were resuspended in RPMI 1640 media (Thermo Fisher) supplemented with 20 U/mL penicillin, 0.02 mg/mL streptomycin, and 10% heat-inactivated FBS. Splenocytes were counted using a Sysmex XT-2000iV haematology analyzer and then used in downstream assays.

### Enzyme-linked immunosorbent spot (ELISpot) assay

IFNγ secretion from splenocytes was measured using a Murine IFNγ Single-Color Enzymatic ELISPOT assay (ImmunoSpot) according to the manufacturer’s protocol. For stimulation, splenocytes were stimulated with 1 µg/mL of an overlapping SARS-CoV-2 Omicron XBB.1.5.X Spike Protein Peptide Pool (StemCell Technologies Cat# 100-1422). DMSO was used as a negative control. Plates were incubated for 20 h at 37°C 5% CO_2_ before being developed. Spots were counted using an ImmunoSpot® S6 Analyzer and reported per million splenocytes.

### Intracellular Cytokine Staining

Splenocytes were stimulated with 1 µg/mL of an overlapping peptide pool (as in ELISpot) for 5 h at 37 °C, 5% CO_2_. Samples were supplemented with GolgiPlug and GolgiStop Protein Transport Inhibitors (BD Bioscience) 45 minutes into the stimulation. Stimulated cells were washed with PBS and then incubated with a viability stain and surface marker antibody cocktail for 30 minutes at 4°C. The cocktail contained LIVE/DEAD™ Fixable Violet Dead Cell Stain (Thermo Fisher), anti-CD3 PerCP-Vio® 700 REAfinity™ (Miltenyi Biotec Cat# 130-120-826, RRID:AB_2752207), anti-CD4 VioGreen™ REAfinity™ (Miltenyi Biotec Cat# 130-118-693, RRID:AB_2734087), and anti-CD8a PE-Vio® 770 REAfinity™ (Miltenyi Biotec Cat# 130-118-946, RRID:AB_2733251). Stained cells were washed with PBS + 0.5% BSA + 2 mM EDTA (FACS buffer) and then fixed/permeabilized using BD cytofix/cytoperm (BD Biosciences) for 20 minutes at 4°C. Cells were washed in Perm/Wash (BD Bioscience) and then stained with intracellular antibody cocktail for 20 minutes at 4°C. This cocktail contained anti-IFNγ FITC REAfinity™ (Miltenyi Biotec Cat# 130-123-283, RRID:AB_2819467), anti-TNF-α PE REAfinity™ (Miltenyi Biotec Cat# 130-119-561, RRID:AB_2784485), and anti-IL-2 APC REAfinity™ (Miltenyi Biotec Cat# 130-129-192, RRID:AB_2922001). Cells were resuspended in FACS buffer and stored at 4°C prior to analysis by flow cytometry (FACSymphoney A1) the next day. Data analysis was completed using FlowJo version 10.10.0 (RRID:SCR_008520). Gating strategy is shown in **Supplementary Figure 4**.

### Quantification and statistical analysis

Statistical analyses were performed using GraphPad Prism 9 (RRID:SCR_002798). A RM two-way analysis of variance (ANOVA) with Tukey’s multiple comparisons test was used for pairwise (between-group) comparisons of body weight data. For all other data, a one-way ANOVA with Tukey’s adjustment was applied for pairwise comparisons of data or log-transformed data. The number of samples for each graph is indicated in the figure legends. In all datasets, ∗p < 0.05, ∗∗p < 0.01, ∗∗∗p < 0.001, ∗∗∗∗p < 0.0001.

## Data Availability

All data supporting the conclusions of this study are present in the main text and supplementary materials. Additional information is available from the corresponding authors upon request.

## Acknowledgements

We gratefully acknowledge the histology and staining services provided by the Louise Pelletier HCF at the University of Ottawa. We gratefully acknowledge the technical contribution of Simon Lord-Dufour, Brian Cass and Louis Bisson at the NRC-HHT for recombinant spike production. We also would like to acknowledge the assistance provided by the Animal Care Facility staff at Health Canada and the National Research Council of Canada. We thank Dr. Lu Huixin and Dr. Roger Tam for commenting on the manuscript and Greg Harris for preparing the photomicrographs. Schematic representations were created in BioRender. This work is supported by the Government of Canada (intramural funding from Health Canada).

## Author contribution statement

Conceptualization, L.T., A.T., M.J.W.J., X.L.; Methodology, L.T., C.L., W.Z., G.F., M.S., A.T.; Formal Analysis and Visualization, L.T., A.S.; Investigation, L.T., C.L., W.Z., D.D., J.B., G.F., A.P., S.N.R. J.W., C.G., A.S., W.C.; Resources, C.L., S.N.R., C.G., M.S., Y.D., A.T.; Writing - Original Draft, L.T.; Writing - Review & Editing, L.T., C.L., W.Z., D.D., J.B., G.F., A.P., S.N.R. J.W., C.G., A.S., M.S., Y.D., W.C., L.W., S.S., A.T., M.J.W.J., X.L.; Supervision, Y.D., L.W., S.S., A.T., M.J.W.J., X.L.; Project Administration L.T., A.T., M.J.W.J., X.L.; Funding Acquisition, Y.D., L.W., S.S., A.T., M.J.W.J., X.L.;

## References

1. Lasrado, N. et al. Waning immunity and IgG4 responses following bivalent mRNA boosting. Sci Adv 10, 9945 (2024).

2. Föhse, K. et al. The impact of BNT162b2 mRNA vaccine on adaptive and innate immune responses. Clinical Immunology 255, 109762 (2023).

3. Ishii, T. et al. Waning cellular immune responses and predictive factors in maintaining cellular immunity against SARS-CoV-2 six months after BNT162b2 mRNA vaccination. Sci Rep 13, 1–12 (2023).

4. Haq, M. A. et al. Antibody longevity and waning following COVID-19 vaccination in a 1-year longitudinal cohort in Bangladesh. Sci Rep 14, 1–11 (2024).

5. Tong, X. et al. Waning and boosting of antibody Fc-effector functions upon SARS-CoV-2 vaccination. Nature Communications 2023 14:1 14, 1–15 (2023).

6. Moore, M., Anderson, L., Schiffer, J. T., Matrajt, L. & Dimitrov, D. Durability of COVID-19 vaccine and infection induced immunity: A systematic review and meta-regression analysis. Vaccine 54, 126966 (2025).

7. Srivastava, K. et al. SARS-CoV-2-infection-and vaccine-induced antibody responses are long lasting with an initial waning phase followed by a stabilization phase. Immunity 57, 587–599.e4 (2024).

8. Widge, A. T. et al. Durability of Responses after SARS-CoV-2 mRNA-1273 Vaccination. New England Journal of Medicine 384, 80–82 (2021).

9. Doria-Rose, N. et al. Antibody Persistence through 6 Months after the Second Dose of mRNA-1273 Vaccine for Covid-19. New England Journal of Medicine 384, 2259–2261 (2021).

10. Pegu, A. et al. Durability of mRNA-1273 vaccine–induced antibodies against SARS-CoV-2 variants. Science (1979) 373, 1372–1377 (2021).

11. Kim, W. et al. Germinal centre-driven maturation of B cell response to mRNA vaccination. Nature 2022 604:7904 604, 141–145 (2022).

12. Nguyen, D. C. et al. SARS-CoV-2-specific plasma cells are not durably established in the bone marrow long-lived compartment after mRNA vaccination. Nat Med 31, 235–244 (2025).

13. Florea, A. et al. Effectiveness of Messenger RNA-1273 Vaccine Booster Against Coronavirus Disease 2019 in Immunocompetent Adults. Clinical Infectious Diseases 76, 252–262 (2023).

14. Abu-Raddad, L. J. et al. Effect of mRNA Vaccine Boosters against SARS-CoV-2 Omicron Infection in Qatar. New England Journal of Medicine 386, 1804–1816 (2022).

15. Ackerson, B. K. et al. Effectiveness and durability of mRNA-1273 BA.4/BA.5 bivalent vaccine (mRNA-1273.222) against SARS-CoV-2 BA.4/BA.5 and XBB sublineages. Hum Vaccin Immunother 20, (2024).

16. Sabnis, S. et al. A Novel Amino Lipid Series for mRNA Delivery: Improved Endosomal Escape and Sustained Pharmacology and Safety in Non-human Primates. Molecular Therapy 26, 1509–1519 (2018).

17. Hassett, K. J. et al. Optimization of Lipid Nanoparticles for Intramuscular Administration of mRNA Vaccines. Mol Ther Nucleic Acids 15, 1–11 (2019).

18. Connors, J. et al. Lipid nanoparticles (LNP) induce activation and maturation of antigen presenting cells in young and aged individuals. Communications Biology 2023 6:1 6, 1–13 (2023).

19. Alameh, M. G. et al. Lipid nanoparticles enhance the efficacy of mRNA and protein subunit vaccines by inducing robust T follicular helper cell and humoral responses. Immunity 54, 2877–2892.e7 (2021).

20. Pfeifle, A. et al. DNA lipid nanoparticle vaccine targeting outer surface protein C affords protection against homologous Borrelia burgdorferi needle challenge in mice. Front Immunol 14, 1020134 (2023).

21. Tursi, N. J. et al. Modulation of lipid nanoparticle-formulated plasmid DNA drives innate immune activation promoting adaptive immunity. Cell Rep Med 6, 102035 (2025).

22. Liao, H. C. et al. Lipid nanoparticle-encapsulated DNA vaccine robustly induce superior immune responses to the mRNA vaccine in Syrian hamsters. Mol Ther Methods Clin Dev 32, 101169 (2024).

23. Guimaraes, L. C. et al. Nanoparticle-based DNA vaccine protects against SARS-CoV-2 variants in female preclinical models. Nature Communications 2024 15:1 15, 1–19 (2024).

24. Li, M. et al. Lipid Nanoparticles Outperform Electroporation in Delivering Therapeutic HPV DNA Vaccines. Vaccines (Basel) 12, 666 (2024).

25. Zhang, W. et al. The Expression Kinetics and Immunogenicity of Lipid Nanoparticles Delivering Plasmid DNA and mRNA in Mice. Vaccines (Basel) 11, 1580 (2023).

26. Patel, M. N. et al. Safer non-viral DNA delivery using lipid nanoparticles loaded with endogenous anti-inflammatory lipids. Nat Biotechnol 1–11 (2025) doi:10.1038/S41587-025-02556-5;SUBJMETA.

27. FDA Takes Action on Updated mRNA COVID-19 Vaccines to Better Protect Against Currently Circulating Variants | FDA. https://www.fda.gov/news-events/press-announcements/fda-takes-action-updated-mrna-covid-19-vaccines-better-protect-against-currently-circulating.

28. Tamming, L. A. et al. DNA Based Vaccine Expressing SARS-CoV-2 Spike-CD40L Fusion Protein Confers Protection Against Challenge in a Syrian Hamster Model. Front Immunol 12, (2022).

29. Tamming, L. et al. Lipid nanoparticle encapsulation of a Delta spike-CD40L DNA vaccine improves effectiveness against Omicron challenge in Syrian hamsters. Mol Ther Methods Clin Dev 32, 101325 (2024).

30. Kaku, Y., Uriu, K., Okumura, K., Ito, J. & Sato, K. Virological characteristics of the SARS-CoV-2 KP.3.1.1 variant. Lancet Infect Dis 24, e609 (2024).

31. Nunes da Silva, W., Dias Moura Prazeres, P. H. & Goulart Guimarães, P. P. PLA-PEG as an alternative to PEGylated lipids for nanoparticle-based DNA vaccination against SARS-CoV-2. Mol Ther Nucleic Acids 35, 102293 (2024).

32. Li, M. et al. Lipid Nanoparticles Outperform Electroporation in Delivering Therapeutic HPV DNA Vaccines. Vaccines (Basel) 12, 666 (2024).

33. Lai, D. C., Nguyen, T. N., Trinh, G. P., Steffen, D. & Vu, H. L. X. Lipid nanoparticle-encapsulated DNA vaccine induces balanced antibody and T-cell responses in pigs with maternally derived antibodies. J Virol 10.1128/JVI.01123-25 (2025) doi:10.1128/JVI.01123-25.

34. Lai, D. C., Nguyen, T. N., Poonsuk, K., McVey, D. S. & Vu, H. L. X. Lipid nanoparticle-encapsulated DNA vaccine encoding African swine fever virus p54 antigen elicits robust immune responses in pigs. Vet Microbiol 305, 110508 (2025).

35. Am, H., et al. CD40 ligand preferentially modulates immune response and enhances protection against influenza virus. J Immunol 193, 722–734 (2014).

36. Kometani, K. & Kurosaki, T. Differentiation and maintenance of long-lived plasma cells. Curr Opin Immunol 33, 64–69 (2015).

37. Pardi, N. et al. Expression kinetics of nucleoside-modified mRNA delivered in lipid nanoparticles to mice by various routes. Journal of Controlled Release 217, 345–351 (2015).

38. Swingle, K. L. et al. Circular RNA lipid nanoparticle vaccine against SARS-CoV-2. Proceedings of the National Academy of Sciences 122, e2505718122 (2025).

39. Qu, L. et al. Circular RNA vaccines against SARS-CoV-2 and emerging variants. Cell 185, 1728–1744.e16 (2022).

40. Wan, J. et al. Circular RNA vaccines with long-term lymph node-targeting delivery stability after lyophilization induce potent and persistent immune responses. mBio 15, (2024).

41. Krawczyk, P. S. et al. Re-adenylation by TENT5A enhances efficacy of SARS-CoV-2 mRNA vaccines. Nature 641, 984–992 (2025).

42. Colwill, K. et al. A scalable serology solution for profiling humoral immune responses to SARS-CoV-2 infection and vaccination. Clin Transl Immunology 11, e1380 (2022).

43. Nie, J. et al. Quantification of SARS-CoV-2 neutralizing antibody by a pseudotyped virus-based assay. Nature Protocols 2020 15:11 15, 3699–3715 (2020).

44. Reed, L. J. & Muench, H. A simple method of estimating fifty per cent endpoints. Am J Epidemiol 27, 493–497 (1938).

45. Wölfel, R. et al. Virological assessment of hospitalized patients with COVID-2019. Nature 2020 581:7809 581, 465–469 (2020).

46. Zivcec, M., Safronetz, D., Haddock, E., Feldmann, H. & Ebihara, H. Validation of assays to monitor immune responses in the Syrian golden hamster (Mesocricetus auratus). J Immunol Methods 368, 24–35 (2011).

47. Livak, K. J. & Schmittgen, T. D. Analysis of Relative Gene Expression Data Using Real-Time Quantitative PCR and the 2−ΔΔCT Method. Methods 25, 402–408 (2001).

48. Harris, G., Holbein, B. E., Zhou, H., Howard Xu, H. & Chen, W. Potential mechanisms of mucin-enhanced acinetobacter baumannii virulence in the mouse model of intraperitoneal infection. Infect Immun 87, (2019).

49. Lien, C. E. et al. CpG-adjuvanted stable prefusion SARS-CoV-2 spike protein protected hamsters from SARS-CoV-2 challenge. Scientific Reports 2021 11:1 11, 1–7 (2021).

