## Supplemental Information for "Durability of DNA-LNP and mRNA-LNP Vaccine-Induced Immunity Against SARS-CoV-2 XBB.1.5"

**Supplementary Information**

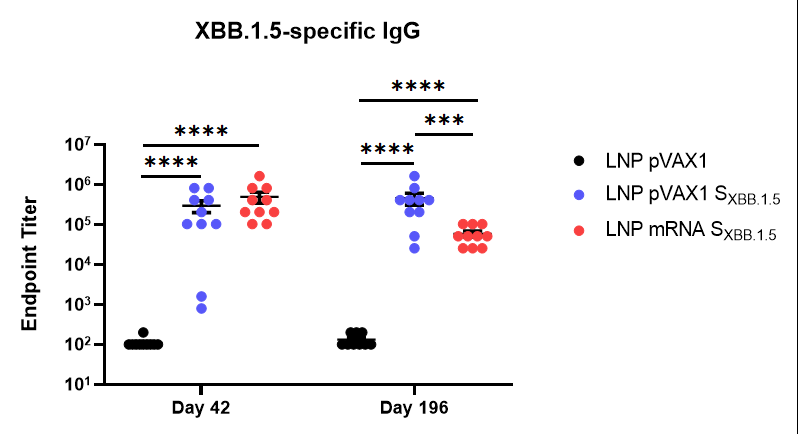

**Supplementary Figure 1. Longevity of DNA-LNP immunity is independent of CD40L.** Male Syrian hamsters were immunized intramuscularly on day 0 and 28 with 5 µg of S_XBB.1.5_ DNA-LNP or 2.5 µg of S_XBB.1.5_ mRNA-LNP. Control hamsters were vaccinated with 5 µg of pVAX1 DNA-LNP. XBB.1.5 Spike-specific IgG in the serum was determined by ELISA on day 42 or 196 post-vaccination. Data shown are mean ± SEM, n= 10. ∗∗∗p < 0.001, ∗∗∗∗p < 0.0001.

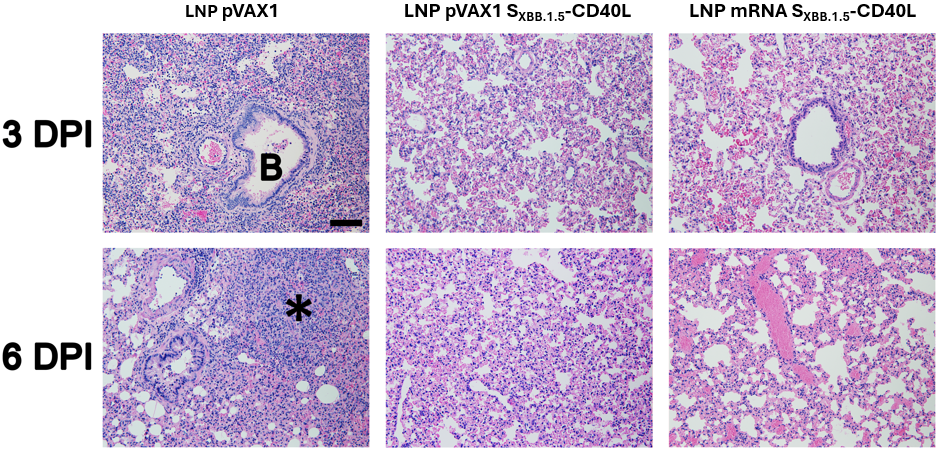

**Supplementary Figure 2. Short-Term Lung Pathology.** Representative photomicrograph of H&E stained lung tissues. Immunized Syrian hamsters were intranasally challenged with an isolate of SARS-CoV-2 XBB.1.5 on day 49. B, bronchioles; *, area of inflammatory cell infiltration and tissue consolidation. Scale bar, 100 μm. Related to Figure 3.

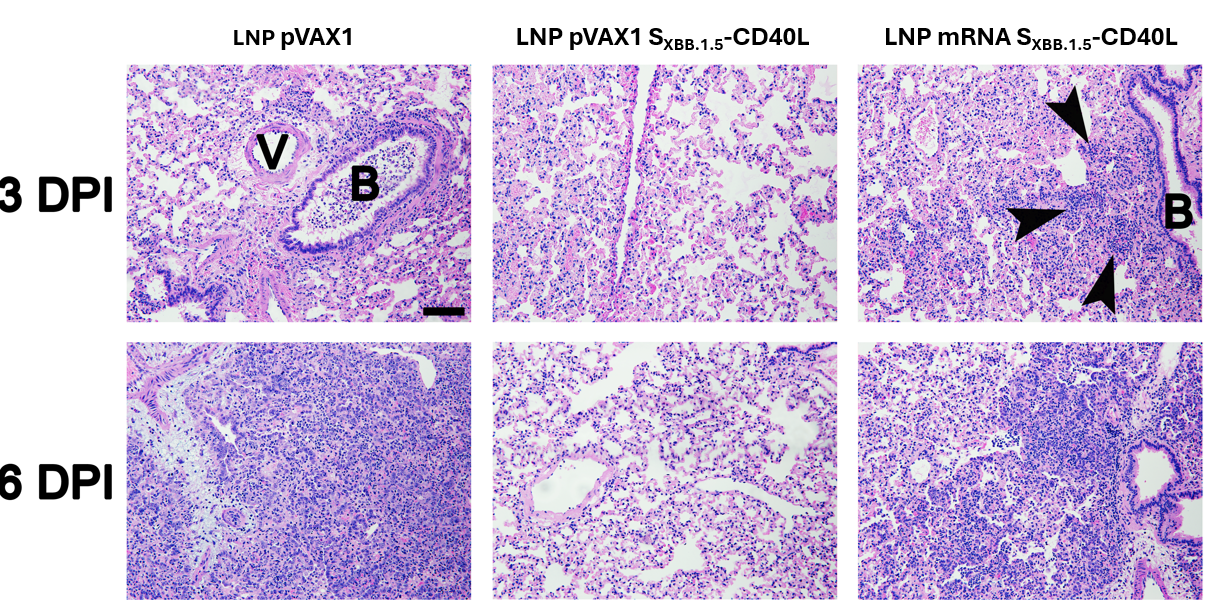

**Supplementary Figure 3. Long-Term Lung Pathology.** Representative photomicrograph of H&E stained lung tissues. Immunized Syrian hamsters were intranasally challenged with an isolate of SARS-CoV-2 XBB.1.5 on day 203. B, exudate of inflammatory cells in the lumen of a bronchus; V, blood vessel. Arrows, peri-airway inflammatory cell infiltration and tissue consolidation. Scale bar, 100 μm. Related to Figure 4.

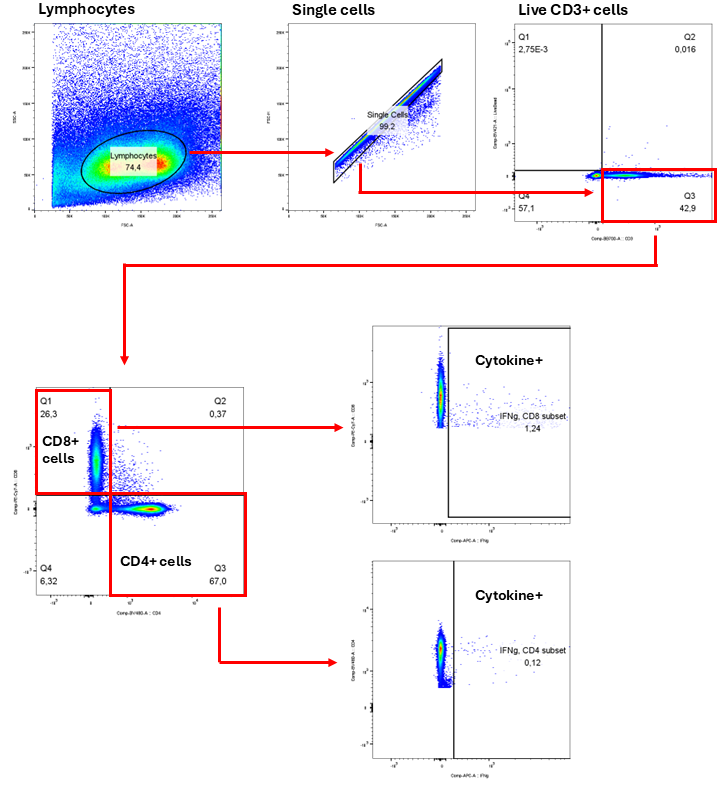

**Supplementary Figure 4. T cell gating strategy.** Gating strategy for cytokine expression in CD4^+^ and CD8^+^ T cells isolated from BALB/c spleens. Representative plots from a S_XBB.1.5_-CD40L mRNA-LNPs vaccinated mouse. Related to Figure 1.

**Supplementary Table 1. Characterization of mouse DNA- and mRNA-LNP vaccines.** Biophysical characterization parameters of DNA- and mRNA-LNP vaccines used in BALB/c studies. Mean particle diameter measured by nanoparticle tracking analysis (NTA). Encapsulation efficiency determined by SYBR™ Gold assay. NT, nucleotides. BP, base pairs. SD, standard deviation. Related to Figure 1.

| **Vaccine**  **(length, NT or BP)** | **Prime/Boost** | **LNP Diameter**  **(Mean ± SD , nm)** | **Encapsulation efficiency (%)** |
| --- | --- | --- | --- |
| pVAX1  (2999) | Prime | 67.5 ± 10.7 | 97.6 |
|  | Boost | 69.1 ± 12.1 | 96.0 |
| mRNA S_XBB.1.5_  (3659) | Prime | 69.2 ± 13.0 | 88.5 |
|  | Boost | 74.0 ± 14.9 | 86.0 |
| mRNA S_XBB.1.5_-CD40L  (4102) | Prime | 72.2 ± 14.8 | 83.9 |
|  | Boost | 75.4 ± 17.6 | 80.7 |

**Supplementary Table 2. Characterization of hamster DNA- and mRNA-LNP vaccines.** Biophysical characterization parameters of DNA- and mRNA-LNP vaccines used in Syrian hamster studies. Mean particle diameter measured by NTA. Encapsulation efficiency determined by SYBR™ Gold assay. NT, nucleotides. BP, base pairs. SD, standard deviation. Related to Figure 2-4 and Supplementary Figure 1.

| **Vaccine**  **(length, NT or BP)** | **Prime/Boost** | **LNP Diameter**  **(Mean ± SD , nm)** | **Encapsulation efficiency (%)** |
| --- | --- | --- | --- |
| pVAX1  (2999) | Prime | 72.2 ± 14.6 | 93.4 |
|  | Boost | 71.3 ± 14.2 | 94.0 |
| pVAX1 S_XBB.1.5_  (6658) | Prime | 79.7 ± 18.2 | 92.4 |
|  | Boost | 78.5 ± 18.7 | 91.8 |
| pVAX1 S_XBB.1.5_-CD40L  (7101) | Prime | 81.3 ± 19.5 | 94.3 |
|  | Boost | 81.7 ± 20.9 | 93.1 |
| mRNA S_XBB.1.5_  (3659) | Prime | 71.6 ± 15.2 | 83.8 |
|  | Boost | 71.7 ± 13.3 | 88.4 |
| mRNA S_XBB.1.5_-CD40L  (4102) | Prime | 72.2 ± 16.7 | 80.7 |
|  | Boost | 72.2 ± 16.1 | 85.2 |
